# PathFinder: Bayesian inference of clone migration histories in cancer

**DOI:** 10.1101/2020.07.10.197194

**Authors:** Sudhir Kumar, Antonia Chroni, Koichiro Tamura, Maxwell Sanderford, Olumide Oladeinde, Vivian Aly, Tracy Vu, Sayaka Miura

## Abstract

**Summary:** Metastases form by dispersal of cancer cells to secondary tissues. They cause a vast majority of cancer morbidity and mortality. Metastatic clones are not medically detected or visible until later stages of cancer development. Thus, clone phylogenies within patients provide a means of tracing the otherwise inaccessible dynamic history of migrations of cancer cells. Here we present a new Bayesian approach, *PathFinder*, for reconstructing the routes of cancer cell migrations. *PathFinder* uses the clone phylogeny and the numbers of mutational differences among clones, along with the information on the presence and absence of observed clones in different primary and metastatic tumors. In the analysis of simulated datasets, *PathFinder* performed well in reconstructing migrations from the primary tumor to new metastases as well as between metastases. However, it was much more challenging to trace migrations from metastases back to primary tumors. We found that a vast majority of errors can be corrected by sampling more clones per tumor and by increasing the number of genetic variants assayed. We also identified situations in which phylogenetic approaches alone are not sufficient to reconstruct migration routes.

**Conclusions:** We anticipate that the use of *PathFinder* will enable a more reliable inference of migration histories, along with their posterior probabilities, which is required to assess the relative preponderance of seeding of new metastasis by clones from primary tumors and/or existing metastases.

**Availability:** PathFinder is available on the web at https://github.com/SayakaMiura/PathFinder.

## 1 Introduction

Metastasis (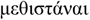, in Gr., to change or transfer) is the spread of abnormal cells from the initiated (the primary tumor) anatomical site to secondary tissues. Cancer is estimated to cause worldwide more than 1.8 million deaths a year (Siegel *et al*., 2020). More than 90% of cancer morbidity and mortality are due to metastases (Welch and Hurst, 2019). Primary tumor cells may seed metastases both locally and at a distance, including different organs, and cells from metastases may also seed new tumors.

Over time, cells in primary tumors and metastases undergo mutations, producing extensive intra- and inter-tumor genetic heterogeneity observed in patients (Williams *et al*., 2019). The genetic variation found in tumors can be used to infer evolutionary relationships of cancer clones within patients as well as migration paths of cancer cells that have seeded and formed metastases (El-Kebir *et al*., 2018; Chroni *et al*., 2019; Alves *et al*., 2019; Miura *et al*., 2018; Somarelli *et al*., 2020). Essentially, the genetic heterogeneity of tumors and cancer clones is becoming a valuable tool to map the origin and progression of cancer in patients. In these efforts, molecular evolutionary and phylogenetic approaches are useful for deciphering how cancer cells evolve, and the pathways of their move from the site of origin to other anatomical sites (Miura *et al*., 2020; Somarelli *et al*., 2017; Alves *et al*., 2019; Chroni *et al*., 2019; El-Kebir *et al*., 2018).

For example, **Figure 1a** shows a phylogeny of five observed clones (C1-C5), and their tumor locations in a patient with colorectal cancer (CRC2 patient) (Leung *et al*., 2017). In this patient, the primary (P) tumor was found in the colon and metastasized to the liver (M). Based on the clone phylogeny and the location of observed clones, Leung et al. (2017) concluded that a polyclonal migration event seeded the metastasis in the colon. That is, multiple genetically different clones from the colon seeded the metastasis. In this example, ancestral clones A3 and A4 are the progenitors of the two clones that seeded the metastasis in the liver (Leung *et al*., 2017).

**Figure 1.**
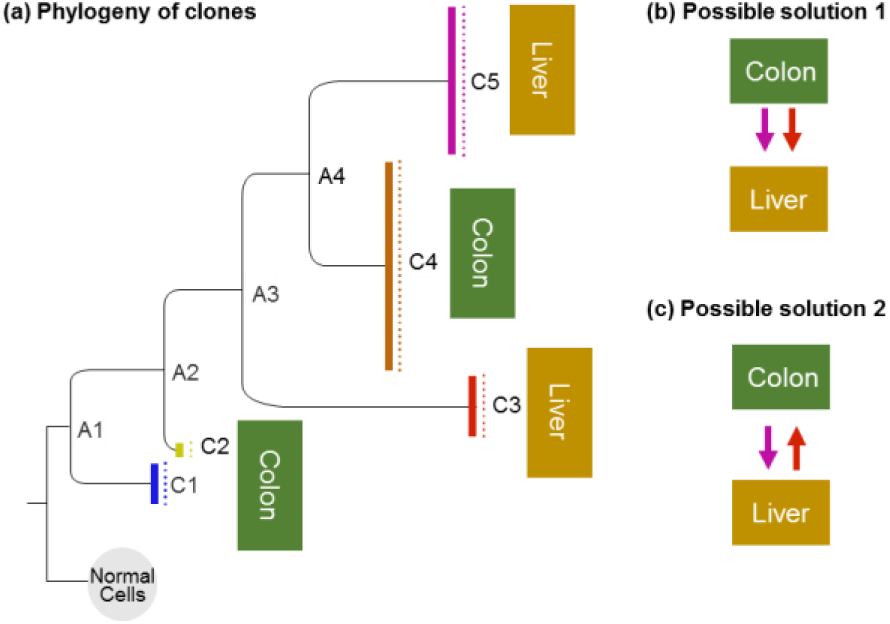
(**a**) A phylogeny of cancer cells in a metastatic colorectal cancer patient (CRC2 patient); redrawn from Zafar et al. (2019). Cancer cells with the same genotype comprise clones (C1-C5), and the lengths of branches are proportional to the number of sequence differences between clones. The phylogeny is rooted on the germline sequence, which represents a healthy and not-mutated cell sequence (Normal). Here, the primary tumor was found in the colon and contained three clones (C1, C2, and C4), whereas the metastatic tumor occurred in the liver and contained two clones (C3 and C5). In addition to the presence of five clones in this patient, this phylogeny shows that at least four other clones existed (ancestral clones, A1 – A4). (**b**) Migration history in which two different clones from the colon, together or at different times, migrated to the liver and seeded metastases. This solution was inferred by Leung et al. (2017) and, further supported by Zafar et al. (2019) who applied the MACHINA approach (El-Kebir et al., 2018). (**c**) An alternative migration history in which clones travelled from colon to liver, but also from liver to colon, after the formation of the metastasis from clones from the primary tumor. This migration history was inferred by MACHINA when the number of tumor sources of seed clones was not constrained (El-Kebir et al., 2018).

Because both Leung et al. (2017) and Zafar et al. (2019) inferred two P→M cell migration paths (**Fig. 1b**), the ancestral clone location (ACL) is estimated to be P for both A3 and A4. Zafar et al. (2019) used MACHINA, a computational approach in which the number of migration events, as well as the number of tumors acting as the source of migration clones, are minimized (El-Kebir *et al*., 2018). In counting the number of migration paths, MACHINA considers multiple cell migrations between the same two tumors (co-migrations) as a single one migration event. The minimization of the number of migration events is equivalent to the maximum parsimony principle in molecular phylogenetics; it is used to infer ancestral states and phylogenetic trees. However, the maximum parsimony approach of MACHINA does not use the information on the amount of genetic differentiation among clones, which varies extensively in clone phylogenies, as seen in **Figure 1a**. Some observed clones show a minimal difference from their ancestral progenitor clones (e.g., C2 from A2), whereas others show much larger differences (e.g., C3 from A3). Therefore, a probabilistic approach is likely to improve the accuracy of migration histories inferred, beyond those made possible by the maximum parsimony approaches.

In this article, we describe a computational method, named as *PathFinder*, that uses not only the evolutionary relationship but also the genetic differentiation among clones to infer migration paths. The importance and significance of a probabilistic approach are evident from the toy example shown in **Figure 2**. When the branch length information is not available, ACL for ancestral clone A2 can be P, M1, or M2, making it impossible to distinguish among the three possible migration histories (**Fig. 2a**.**I-III**). However, when observing the clone phylogeny with branch lengths, we see that ancestral A2 and observed C2 clones are genetically identical. So, one would intuitively infer that A2 is found in the same tumor as does the observed clone C2, i.e., ACL for A2 is likely M1 (**Fig. 2b**).

**Figure 2.**
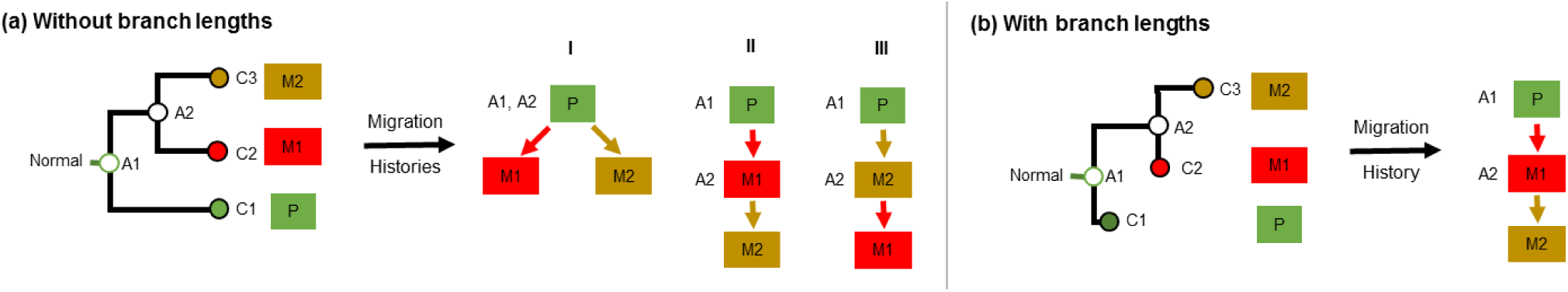
Clone phylogeny (a) without branch lengths and (b) with branch lengths. In panel a, three possible migration histories are shown, because the ancestral clone A2 may have been present in the primary tumor or in one of the two metastases. In panel b, the most likely migration history is shown based on the clone phylogeny with branch lengths, because A2 is nearly identical to clone C2 (and genetically different from clones C1 and C3). Branch lengths provide crucial information. The information deducted by branch lengths can be used into giving insight for choosing the most likely migration path (P→M1→M2).

Consequently, the most likely migration history is P→M1→M2. The *PathFinder* approach, described in the next section, predicts that the P→M1→M2 path is much more likely than the other two possibilities. In contrast, MACHINA infers independent seedings of the two metastases from the primary tumor (P→M1 and P→M2) as the most probable migration scenario. This is because MACHINA does not use branch lengths and minimizes the number of sources that contribute seed clones. Therefore, the inference of the origin and movements of tumor clones will benefit from the use of a probabilistic approach.

*PathFinder* employs a Bayesian statistical molecular phylogenetic framework for inferring ancestral states (ACLs) and generates clone migration pathways between tumors that have the highest posterior probabilities (PPs). *PathFinder*’s probabilistic approach enables us to select from alternative hypotheses of clone migrations statistically. For example, *PathFinder* will allow one to distinguish between the polyclonal seeding and reseeding events (**Fig. 1b** and **1c**, respectively) as well as the source of seeding of new tumors, i.e., primary tumor versus metastasis (e.g., **Fig. 2**). Such distinctions are essential for our understanding of metastasis. It is now becoming clear that metastatic processes are complex with multiple clones seeding tumors, multiple tumors acting as the source of migrations, and even bidirectional seeding events occurring (Sanborn *et al*., 2015; Choi *et al*., 2017; Hoadley *et al*., 2016; Gundem *et al*., 2015; Eirew *et al*., 2015; Brown *et al*., 2017)

In the following, we present the *PathFinder* approach. Then, we show its accuracy in inferring metastatic migration histories by using computer-simulated datasets in which metastases were seeded by only single clones (monoclonal) or by multiple clones (polyclonal), and seeding sources included metastases, in addition to primary tumors. We compare the performance of *PathFinder* with MACHINA. We also assessed the impact of minimization of tumor-sources and preference of co-migration pathways, which are used in MACHINA, on *PathFinder*’s probabilistic inferences (El-Kebir *et al*., 2018). Finally, we applied *PathFinder* in the analysis of datasets from patients with basal-like breast cancer to show its utility (Hoadley *et al*., 2016).

## 2 Methods

### 2.1 The *PathFinder* Method

*PathFinder* assumes that the clone phylogeny, the alignment of clone sequences, and the anatomical locations of every observed clone are known. Using this information, *PathFinder* will infer the location for every ancestral clone (ACL) by using a Bayesian approach and build clone migration histories. For simplicity, we use a phylogeny containing three clones (C_1_, C_2_, and C_3_) that are found in a primary tumor (P) and two metastases (M1 and M2), respectively (**Fig. 3**). In this phylogeny, the normal cells serve as the outgroup, and there are two ancestral clones (A_1_ and A_2_) for which the anatomical location is not known. *PathFinder* infers ACLs for A_1_ and A_2_ by advancing the Bayesian approach of ancestral state inference (Yang *et al*., 1995) (**Fig. 3**). In this case, we estimate branch lengths of the clone phylogeny by using the clone sequence alignment along with the estimates of ACLs. In this joint inference, the instantaneous rates of state changes between the presence and absence of variants are assumed to be equal, and between different tumor states are assumed to be equal as well.

In this case, *x*_1_, *x*_2_, and *x*_3_ represent the location of clones C_1_, C_2_, and C_3_, respectively. *y*_1_ and *y*_2_ are the ACLs of clones A_1_ and A_2_. Vector ***x*** = (*x*_1_, *x*_2_, *x*_3_) and ***y*** = (*y*_1_, *y*_2_). The probability of observing a given configuration of ***x*** is

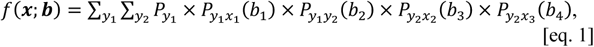

where ***b*** = (*b*_1_, *b*_2_, *b*_3_, *b*_4_) is the vector of branch lengths in the example clone phylogeny derived from clone sequence alignment. Here, *P*_*ij*_(*b*_*k*_) is the probability that the given clone will remain in the same location (*i* = *j*) or move to a different location (*i* ≠ j) after *b*_*k*_ substitutions on branch *k*. To compute *P*_*ij*_(*b*_*k*_), we use a mathematical model of instantaneous state change in which the probability of movement from any locations to another location is equal.

Pursuing the Bayesian approach for computing the posterior probability of each possible configuration for two ancestral clones ***y*** = (*y*_1_, *y*_2_), we write:

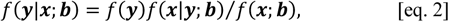

where *f*(***y***) is the prior probability of occurrence of ***y*** and is given by

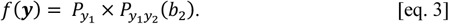

The conditional probability of observing ***x*** for a given set of ancestral clone locations ***y*** is:

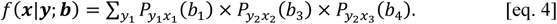

Using this information, we compute the posterior probability of the presence of an ancestral clone (e.g., A_2_) in the metastasis M1 by

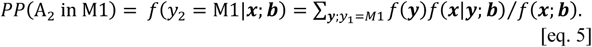

**Figure 3.**
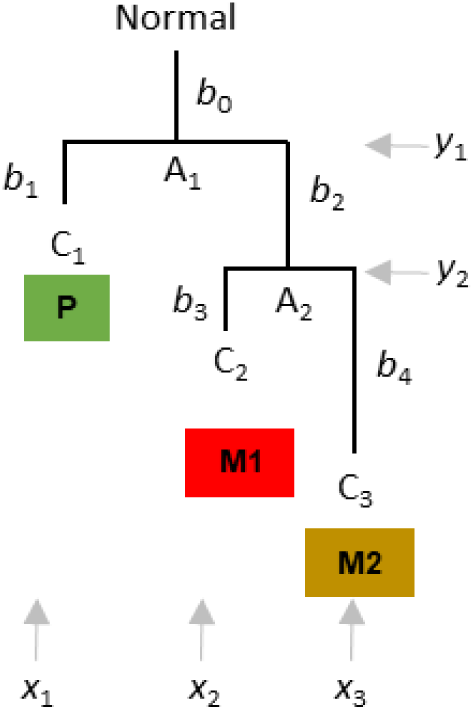
A phylogeny of three clones (C1, C2, and C3) found in three tumors (P, M1, and M2). Clone relationships with branch lengths (*b*’s) are shown, along with the locality in which each clone is found. A1 and A2 are the ancestral clones, and “Normal” refers to the germline/non-cancer cell sequence.

Similarly, we compute the posterior probability of the presence of A_2_ in metastasis M2 and primary tumor P. The ACL for A_1_ will then be the location with the highest posterior probability. By default, *PathFinder* assumes that the seeding events began from the primary tumor, e.g., (El-Kebir *et al*., 2018), so we set ACL(A_1_) = P.

In the explanation above, for simplicity purposes, each clone was assumed to be present in only one location. However, in tumor datasets from patients, we often encounter the same clone in multiple locations. For these datasets, we include each such clone in the clone phylogeny as many times as the number of different locations in which it is present. We append a data column to the clone sequence alignment, which contains the tumor location. In this way, tips of the clone phylogeny are distinguished by their location in the phylogeny used in *PathFinder*.

After estimating PP of all ACLs for all the ancestral clones in the clone phylogeny, we traverse the clone tree to generate all possible migration histories (MH) as directed graphs of cell migrations whenever ACLs are not the same for the pair of nodes connected by a branch. The probability of migration history, *P*_MH_, is simply the product of the posterior probabilities of all the ACLs involved in that history. By default, the graph with the highest *P*_MH_ is chosen to represent the migration history. By the way, if a clone phylogeny is not strictly bifurcating, i.e., some nodes give rise to more than two descendants, then *PathFinder* will explore all possible sets of candidate bifurcations for each polytomy (e.g., three alternative bifurcations for a polytomy involving three branches) to select the ACL that receives the highest PP by applying equations 1-5 to alternative phylogenies.

Alternatively, one may sum the probability of each cell migration edge over all possible migration histories and then assemble a consensus migration history (cMH). One may specify a threshold PP to consider an ACL to be included in the candidate list for generating a collection of possible MHs; we used a cut-off of 0.15, but similar results were obtained by using 0.05. In the final reconstruction, the researcher has the option only to retain migration edges that showed an edge probability of 0.5 or higher, which we found to be very effective in removing spurious edges. With this option, we found that maximum probability MH was as accurate as cMH (see **Fig. 8**). It is worth noting that *PathFinder* reports all the alternative migration histories and their normalized probabilities such that the sum of probabilities from all alternative migration histories considered is 1. The software is programmed in python and available for use on Windows machines (https://github.com/SayakaMiura/PathFinder); a Linux version is being developed.

### 2.2 Assembly and Analysis of Computer Simulated Data

To evaluate and benchmark the performance of *PathFinder*, we used an independently available data collection that has been analyzed in multiple studies (El-Kebir *et al*., 2018; Chroni *et al*., 2019). This collection consists of datasets simulated with clone evolution and tumor growth models under various scenarios. For these datasets, the number of tumors sampled varied from 5 to 7, which we refer to as *t*5 datasets, and between 8 and 11, which we refer to as 8-tumor datasets *t*8 dataset. Overall, the number of tumors was 5–11, the number of clones was 6–26, and the number of single nucleotide variants (SNVs) was 9–99 (El-Kebir *et al*., 2018).

The complexity of the simulated datasets varied based on the number of tumor clones migrating, the number of tumor sites acting as sources and/or recipients of migration, and the number of metastatic clones migrating back to the primary tumor. In total, we tested *PathFinder* on 80 simulated datasets and four seeding scenarios determining the complexity of migration paths (**Fig. 4**). The datasets are available from https://github.com/raphael-group/machina. More details about these datasets can be found in El-Kebir et al. (2018) and Chroni et al. (2019).

**Figure 4.**
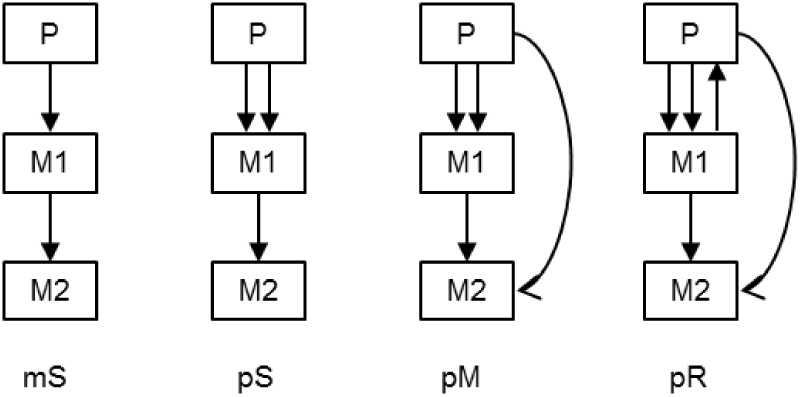
Examples of clone seeding scenarios used for generating simulated data (El-Kebir et al., 2018), arranged by complexity: single clones migrating from single tumor-sources (mS, monoclonal single-source seeding) or from multiple tumors (pS:, polyclonal single-source seeding), and multiple clones migrating from multiple sources (pM, polyclonal multi-source seeding) or migrating from metastasis back to primary (pR, polyclonal reseeding). Redrawn from Chroni et al. (2019).

*PathFinder* software was used to analyze these datasets to generate consensus migration histories (cMH) using the options noted above. For comparative analysis, we retrieved MACHINA results from the PMH-con approach applied by Chroni et al. (2019). PMH-con was chosen because it showed the highest accuracy when compared to PMH-TR and a Bayesian biogeographic approach (BBM) (Chroni *et al*. 2019). Settings in PMH-con included constrained of the primary tumor at the root of the tree, and no restrictions were placed on the possible seeding scenarios and the number of migrations and comigrations.

In all of these analyses, similar to Chroni et al. (2019) approach, the focus was on the accuracy of inference of migration histories when the clone sequences and phylogeny are already known. Errors are usually involved in de-convoluting clones from bulk sequencing data, and in imputing missing data and correcting false positives and false negatives in single-cell sequences exist. However, an analysis of those errors is beyond the scope of our article.

### 2.3 Accuracy measurements

For each migration history, we recorded migration paths inferred correctly (true positives; TPs), migration paths not found (false negative; FNs), and incorrect migration paths (false positives; FPs). We then computed F_1_-score for each dataset, which is the harmonic mean of precision and recall:

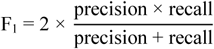

where

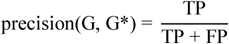

and

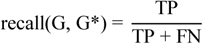

F_1_’s were estimated for individual migration histories inferred. When multiple migration histories were inferred for a dataset, F_1_ represented the simple average of F_1_’s of each migration history. For a collection of datasets, the average F_1_ was also the arithmetic mean of the dataset-specific F_1_’s.

### 2.4 Analysis of an Empirical Data Set

We applied *PathFinder* to A1 and A7 datasets from two patients with basal-like breast cancer (Hoadley *et al*., 2016). The A1 dataset included eight clones from a primary and four metastases (329 SNVs), whereas the A7 dataset consisted of ten clones from primary and five metastases (478 SNVs) (Hoadley *et al*., 2016). We used clone phylogenies that were rooted using the germline sequences (normal cells) as outgroups. We conducted a tumor migration inference analysis in *PathFinder*.

## 3 Results and Discussion

### 3.1 Single-source, monoclonal seeding (*mS*)

The monoclonal (*m*) seeding of metastases represents the simplest scenario of migration histories. In this case, each metastasis was seeded by only one clone, and it received clones from only one tumor (single source, *S*). First, we analyzed the *t*5 datasets consisting of 5-7 tumors. *PathFinder* produced correct migration histories for 9 out of 10 datasets (average F_1_ = 0.975). There was only one error in one dataset in which a P→M3 seeding event was predicted, instead of P→M1→M3. We found this error to be due to insufficient sampling of clones that were present in M1 (**Fig. 5**). The missing clone originated in the primary tumor and was the ancestor of the clone that seeded tumor M3. Therefore, more extensive sampling of clones from each anatomical site would be needed to eliminate such errors.

**Figure 5.**
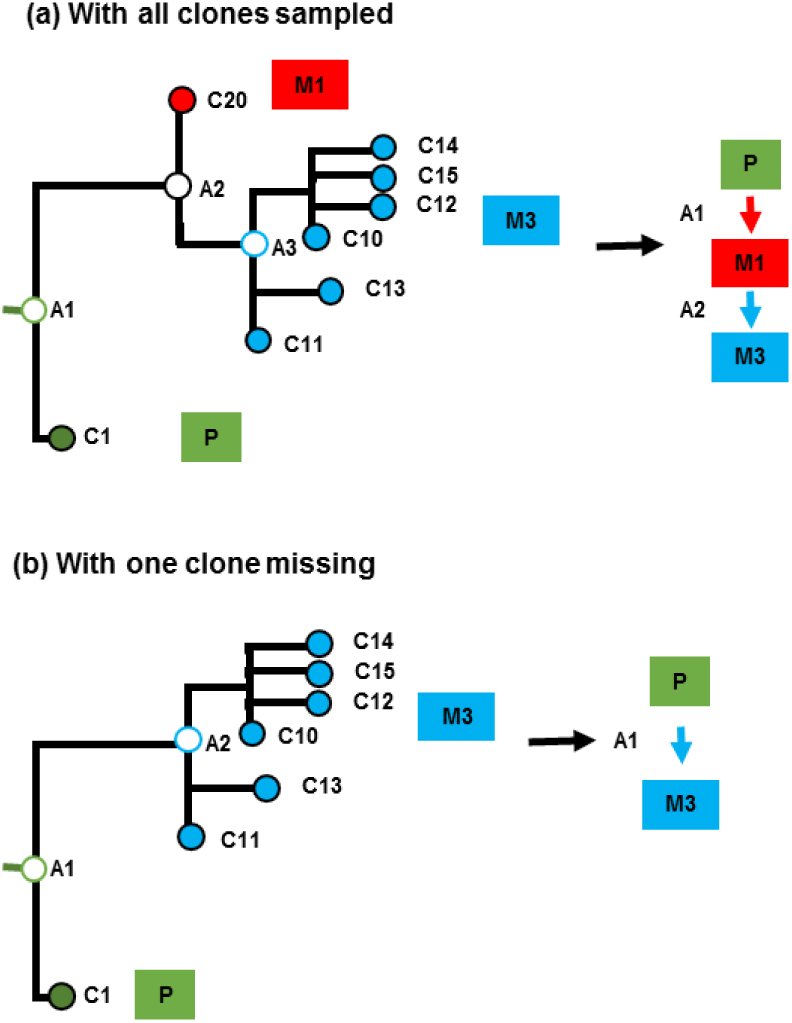
Incomplete clone sampling causes multiple errors. The clone that traveled from M1 to M3 (panel **a**) was not sampled, so a P→M3 seeding event was inferred (false positive), instead of a P→M1 (false negative) and a M1→M3 (false negative) (panel **b**). These three errors cannot be corrected by any computational methods, because there is no way to assess the presence of the ancestral clone A2 without a branching point in the clone phylogeny.

In the *t*8 datasets, there was an increase in the number of tumors (8-11), the number of clones (19.2), as well as the size of the migration histories. The average number of migration paths for these datasets was 7.6, as compared to 4.1 for *t5* datasets. For these datasets, *PathFinder* produced correct migration histories for seven datasets (average F_1_ = 0.92). In one of the three datasets for which *PathFinder* MH contained errors, the problem was caused by the non-sampling of some key primary tumor clones. This problem can only be remedied by sampling more clones per anatomical site.

For the other two datasets, *PathFinder* errors were unavoidable because different clones with identical sequences existed in two source tumors, making it impossible for any computational method to distinguish which tumor provided the seed clones. In practical data analysis, it may be possible to mitigate such errors by sampling more genomic sites (SNPs) that can distinguish clones.

These results show that the inference of migration histories of a large number of anatomical sites increases the complexity of migration paths and requires more extensive clone sampling and the number of SNPs.

### 3.2 Single-source, polyclonal seeding (*pS*)

Next, we present results from the analysis of datasets in which multiple clones seeded metastasis, polyclonal seeding (*p*). However, all the seeding clones came from the same source (*S*) for a given metastasis (of course, different sources may seed different metastasis). In the simulated data, 2-3 clones seeded each metastasis. *PathFinder* produced correct results for eight of the *t*5 datasets (F_1_ = 0.95). In the *t*8 datasets, we observed errors in four migration histories (F_1_ = 0.95). The average number of migration paths for these datasets was 9.1 as compared to 5.5 for *t*5 datasets. For two of the *t*8 datasets, computational errors were unavoidable because of incomplete sampling of clones. The error in the third dataset was caused by the fact that the ancestral clone A2 was equally different from its descendant clones found in two tumors (M1 and M4). In this case, *PathFinder*’s probabilistic approach predicted the ACL to M1 or M4 with similar probabilities, resulting in two equally likely possibilities: P→M4→M1 and P→M1→M4 (**Fig. 6**). For such data, the phylogeny alone is not sufficient, and we need additional information (e.g., mutational signatures, (Christensen *et al*., 2020)) to remedy the lack of resolution.

**Figure 6.**
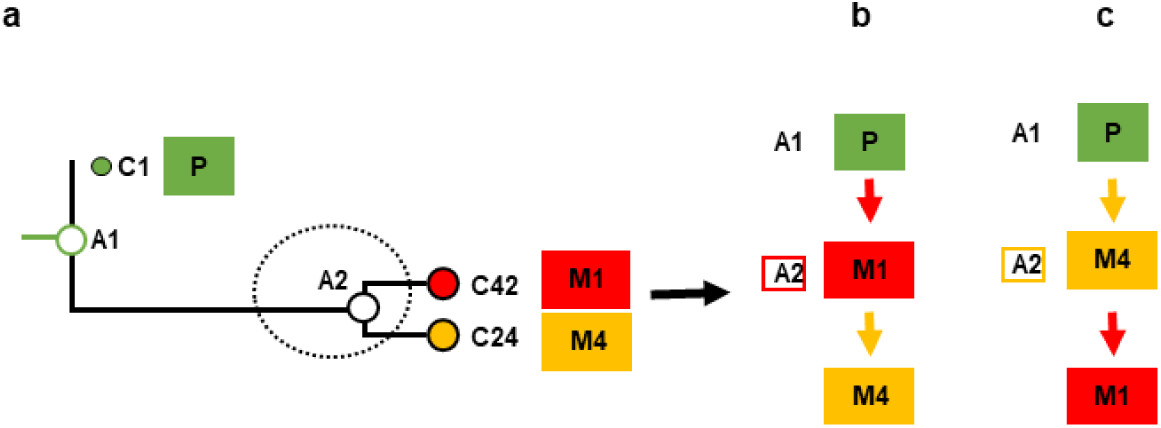
An example from polyclonal seeding scenario in which the migration history inferred by *PathFinder* was inconclusive. Here, the error was caused by the fact that an ancestral clone (A2) is equally different from its descendant clones in tumors M1 and M4, e.g., branch lengths for C24 and C42 clones are very similar (panel **a**). This results in very similar posterior probabilities for two migration paths: P→M4→M1 (panel **b**), and P→M1→M4 (panel **c**). To overcome these type of errors, we need other information (e.g., mutational signatures) in addition to clone phylogenies.

### 3.3 Multiple-source, polyclonal seeding (*pM*)

Next, we explored even more complex and realistic migration histories, in which each metastasis was seeded by multiple clones (2-3, polyclonal *p*) that came from multiple tumors (*M*). For the *t*5 datasets, *PathFinder* predicted correct migration histories for 50% of the datasets (F_1_ = 0.92), with errors found in other datasets again caused by an incomplete sampling of clones, making it computationally impossible to reconstruct some of the migrations. Therefore, a more accurate migration history inference would require more extensive sampling of clones from each tumor.

The migration histories for *t*8 datasets became much more extensive, containing 10.5 migation paths, as compared to 6.8 for *t*5 datasets. The F_1_ score was 1.0 for four of the datasets, 0.90 - 0.96 for four datasets, and lower for the remaining two datasets (0.55 and 0.73). In these cases, again, most of the *PathFinder* errors were due to a lack of sufficient sampling of clones required to detect migrations. Also, many key clones sampled from multiple tumors were identical that made it difficult to discern the origins of seeding clones in the worse performing dataset. Therefore, more extensive sampling of clones and sequencing of additional SNPs will be needed to improve the performance of computational methods.

### 3.4 Multiple-source, polyclonal seeding and reseeding (*pR*)

The most challenging datasets for *PathFinder* were those in which the primary tumor was receiving clones back from one or more metastases. That is, clones migrated from some metastases back to the primary tumor (re-seeding events). These pR datasets were also multiple-source (more than two tumors). They included multiple (1-2) clones seeding metastases and the reseeding events (single or multiple seeding events from metastasis back to primary). For the *t*5 datasets, 60% of the migration paths were entirely correct (F_1_ = 0.89). In the worst-performing dataset, no seeding clones were part of the randomly selected clone sample in one of the source tumors, which meant that one could never infer a vast majority of M→M seeding events as well as reseeding. The *t*8 datasets (F_1_ = 0.75) had an average number of migration paths of 10.1 as opposed to 7.2 for *t*5 datasets. Datasets with incorrect migration graphs presented similar issues as the datasets from the most straightforward seeding scenarios.

### 3.5 Performance by the number of tumors and migrations types

With the sampling of a higher number of tumors (5-7 vs. 8-11), a higher number of clones were also sampled from 13.4 to 20 (a 66% increase, on average), but the error (=1-F_1_) increased from 0.09 to 0.16. This increase is proportional to the increase in the number of migration events that increased by 63%. Therefore, the higher the number of migration events, the more the error in inferring them correctly. Overall, the highest accuracy decrease was seen for simulated datasets that involved reseeding. In these cases, the error increased from 14% to 31% for *t*5 and *t*8 datasets.

As expected, less complex migration histories (*mS* type) were much easier to infer than the complex ones (*pR* type). The overall accuracy of mS, pS, pM, and pR histories are shown in **Figure 7**. This patterns arose because P→M migrations were the easiest to infer (F_1_ = 0.92) followed by M→M (F_1_ = 0.84). F_1_ for M→P was more complex, because *PathFinder* predicted no correct M→P paths for 21 datasets. For others, F_1_ was 1.0. The *mS* migration histories consisted of a lot of P→M migrations (81%), along with a small fraction of M→M migrations (19%). In contrast, *pS, pM*, and *pR* contained many fewer P→M migrations (77%, 64%, and 48%, respectively).

**Figure 7.**
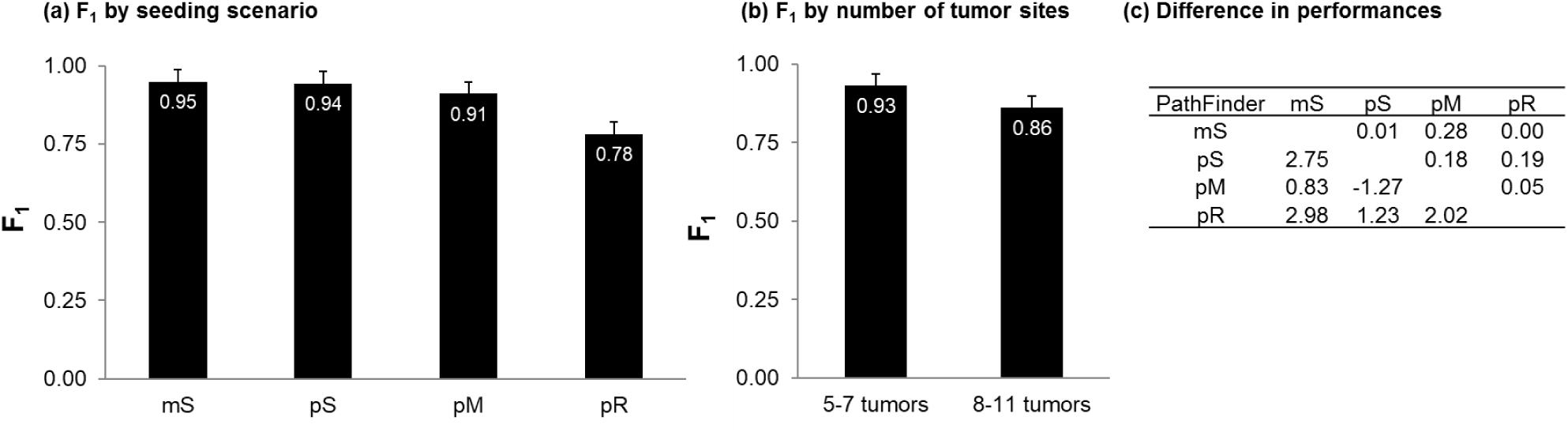
Overall performance of *PathFinder* for different types of (**a**) migration histories and (**b**) datasets with small and large number of tumor sites sampled. Standard errors are also shown. (C) A tabular comparison of the difference in F1-scores of *PathFinder* between the seeding scenarios is shown. The z-scores and the corresponding *P* values are shown below and above the diagonal, respectively.

### 3.6 The usefulness of MACHINA criteria and the performance of most probable MHs

The parsimony approach in MACHINA uses a hierarchical minimization scheme which not only strives to generate the most parsimonious migration history by minimizing the number of migrations but also optimizes the number of co-migrations such that each co-migration is considered a single event. Thus, co-migrations, meaning the events of multiple clones migrating, are preferred. Finally, it minimizes the number of tumors that can act as the sources of seeding clones. We tested if this type of multi-level optimality scheme will be beneficial in selecting more accurate migration histories when *PathFinder* produces multipole MHs with non-zero probability.

There were 31 (out of 80) datasets for which *PathFinder* detected multiple migration histories with different F_1_ scores (0.07 < *P* ≤ 0.81). After applying MACHINA’s hierarchical optimality scheme, the F_1_-scores of the migration histories inferred did not improve in most of the cases (**Fig. 8**). Overall, the average F_1_ for the *PathFinder*’s consensus MH was 0.83, which is close to that after applying MACHINA’s scheme. This could be taken to mean that the use of a probabilistic approach obviates the need to impose a parsimony principle to infer or fine-tune MH inferences. **Figure 8** also shows that the difference between the choice of a consensus MH and the one with the highest probability is rather small, as their F_1_ scores were the same (0.83, respectively). So, one may choose to infer either a consensus or the highest probability migration history in biological analyses.

**Figure 8.**
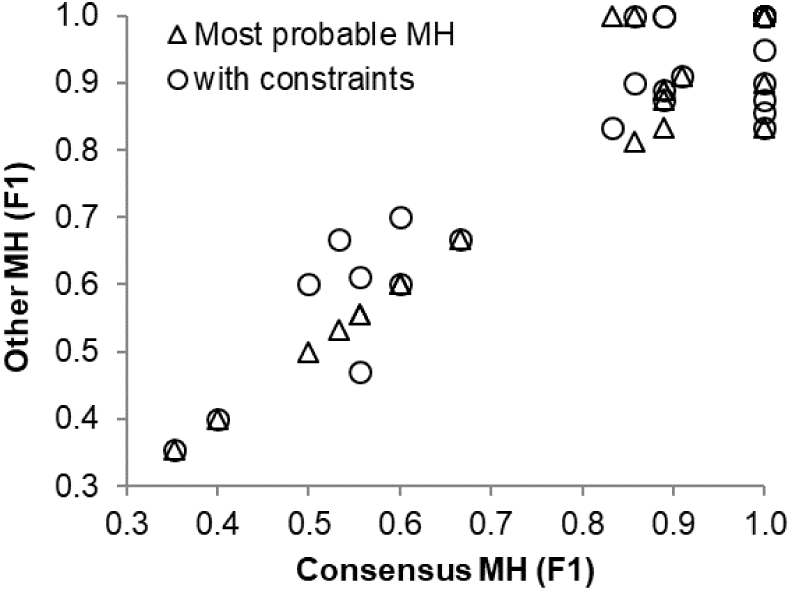
Scatter plot of F1-scores of *PathFinder* (consensus migration history, MH) and (i) of those with MACHINA’s hierarchical (circles) and (ii) of those with *PathFinder*’s most probable MH (triangles). The graph shows the results for which *PathFinder* found multiple alternative MH (31 datasets).

### 3.7 Comparisons of *PathFinder* with MACHINA

In **Figure 9**, we show the comparative performance of *PathFinder* and MACHINA approaches. Results for MACHINA were obtained from Chroni et al. (2019), who also analyzed the same datasets under the same conditions. For simpler cases that involved single clones migrating from single tumors (*mS*), *PathFinder* improved upon MACHINA by 2%. For datasets with polyclonal seeding (*pS*), *PathFinder* improved the performance by 9%. In both cases, as noted earlier, many errors are due to insufficient tumor or SNV sampling, so it is unlikely that one could improve this accuracy much more for these two datasets. The same is likely true for *pM* datasets, in which *PathFinder* performed 10% better than MACHINA. These are significant improvements considering that only 7 – 19% of migration paths were incorrect for these three types of migrations. For the *pR* datasets, with reseeding, which are the most complicated migration histories, did not see a noticeable difference between *PathFinder* and MACHINA (**Fig. 9a**). Finally, *PathFinder* performed better than MACHINA for datasets with small as well as a large number of tumors (**Fig. 9b**). Many of the differences were not statistically significant in the t-tests, mainly because of small sample sizes as the number of datasets analyzed is small within categories.

**Figure 9.**
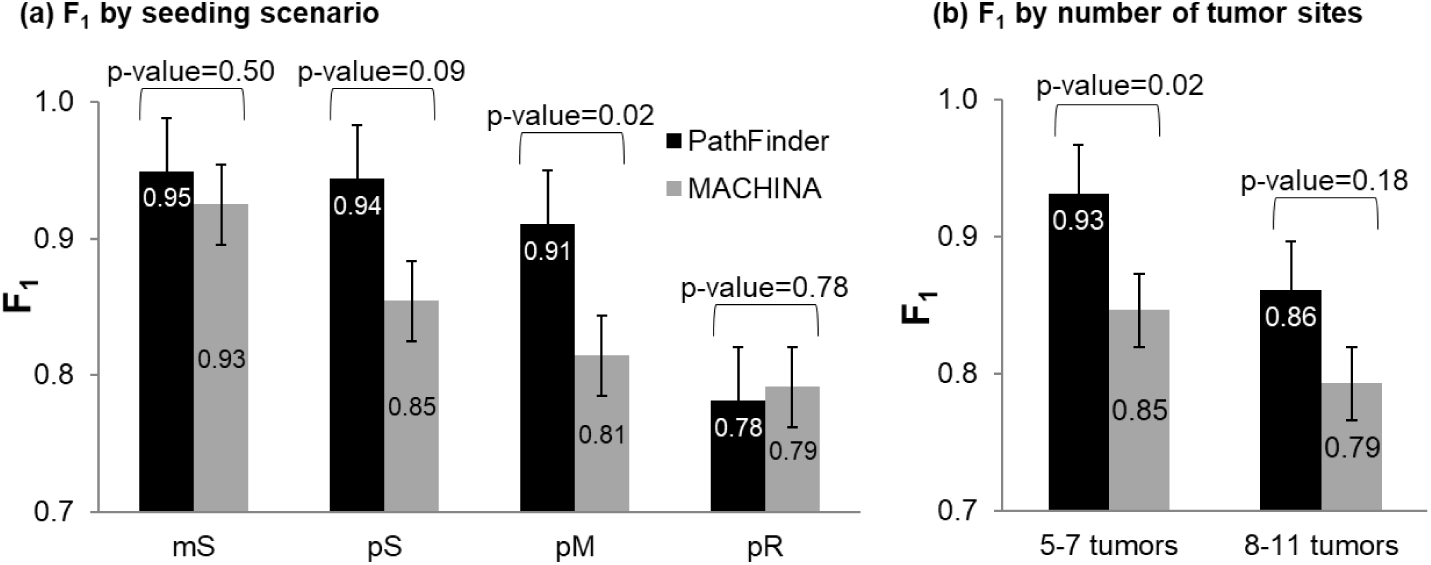
Comparative performance of *PathFinder* (black bars) and MACHINA (gray bars) for (**a**) different types of migration histories and (**b**) datasets with small and large number of tumor sites sampled. Standard errors and p-values by t-test are also shown.

These results establish the utility of a probabilistic (*PathFinder*) over a parsimony based approach (MACHINA), as the clone migration inferences benefited from the use of branch lengths, showing the power of an evolutionary-aware framework on deciphering especially difficult cases with multiple clones moving between tumors. At the same time, it is prudent to acknowledge that even a parsimony based approach, as developed in MACHINA, is adequate for many datasets.

### 3.8 Breast cancer analysis

We applied *PathFinder* to two published datasets of basal-like breast cancer (Patients A1 and A7) (Hoadley *et al*., 2016). We first discuss the A7 dataset that contained ten clones from a primary tumor (breast) and five metastases (brain, lung, rib, liver, and kidney). The evolutionary relationships of these clones and the associated tumor sites are shown in **Figure 10a**. Analyses of these data by *PathFinder* predicted two migration events from primary to metastases (P→M paths) and five migration events between metastases (M→M paths) (**Fig. 10b**). All migration events were highly supported in the Bayesian analyses (*P* = 1.0).

**Figure 10.**
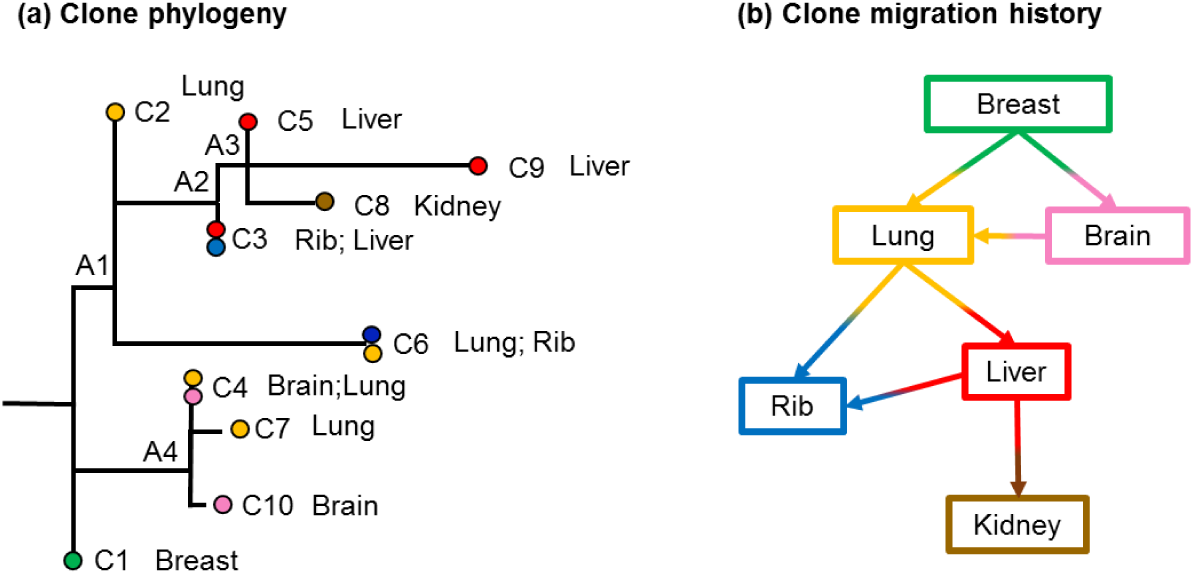
Analysis of Patient A7 with basal-like breast cancer (Hoadley *et al*., 2016). (a) Clone phylogeny and tumor location of each clone reported in the original study. Nodes A1-A4 are ancestral nodes. (b) Clone migration history predicted by *PathFinder*. Inferences of migration paths includes many M→M paths, all of which have a high *P = 1*.*0*. Colors correspond to the tumor location where clones were sampled from.

The P→M paths involved seeding events from the breast tumor to lung and brain metastases (**Fig. 10b**; *P* = 1.0). The breast to lung seeding is inferred because the ancestral clone A1 was estimated to be present in the lung tumor with a *P* = 1.0, because the genetic sequence of observed clone C2 is predicted to be the same as that of A1 (**Fig. 10a**). The brain metastasis is also predicted to be seeded by clones from the breast, the clones found within the brain (C4) are closer to the ancestral clone A4 than the lung clones.

*PathFinder* predicted multiple instances of metastasis to metastasis (M→M) in this patient (**Fig. 10b**). This seeding scenario is different from the conclusion of Hoadley et al. (2016), who proposed that the primary tumor directly seeded all the metastases. We argue that this is not reasonable based on the observed clone phylogeny and the genetic differences between the clones. We explain it, for example, for the cluster containing clones C3, C5, C8, and C9. All of these clones are found in the liver, kidney, and rib metastases. If the migration history proposed by Hoadley et al. (2016) were to be accurate, i.e., breast seeded the metastases in the liver, kidney, and rib, and we would expect some breast tumor clones to be present near their most recent common ancestor (ancestral clones A2 and A3). However, no such clones were observed in the phylogeny, and the best inference in the absence of additional clone sampling is to posit many seedings between metastases. Overall, *PathFinder* suggests many more M→M seedings than the P→M seedings in this patient.

Next, we present the migration history for Patient A1 inferred by *PathFinder*. The A1 dataset included five clones from a primary tumor (breast) and four metastases (adrenal, lung, spinal, and liver) (**Fig. 11a**). *PathFinder* inferred seven P→M, and one M→P paths. There were four more migration events (colored in gray, **Fig. 11b**), but because they were supported by low values of probabilities (<0.5). For this dataset, *PathFinder* predicted that all metastases were founded by clones that migrated from the primary tumor, which is evident from the structure of the phylogeny and consistent with the Hoadley et al.’s conclusions. The ancestral tumor sites for the nodes A1, A3, and A4 were predicted to be the primary tumor, even though the migration inferences are not highly supported (**Fig. 11b**).

**Figure 11.**
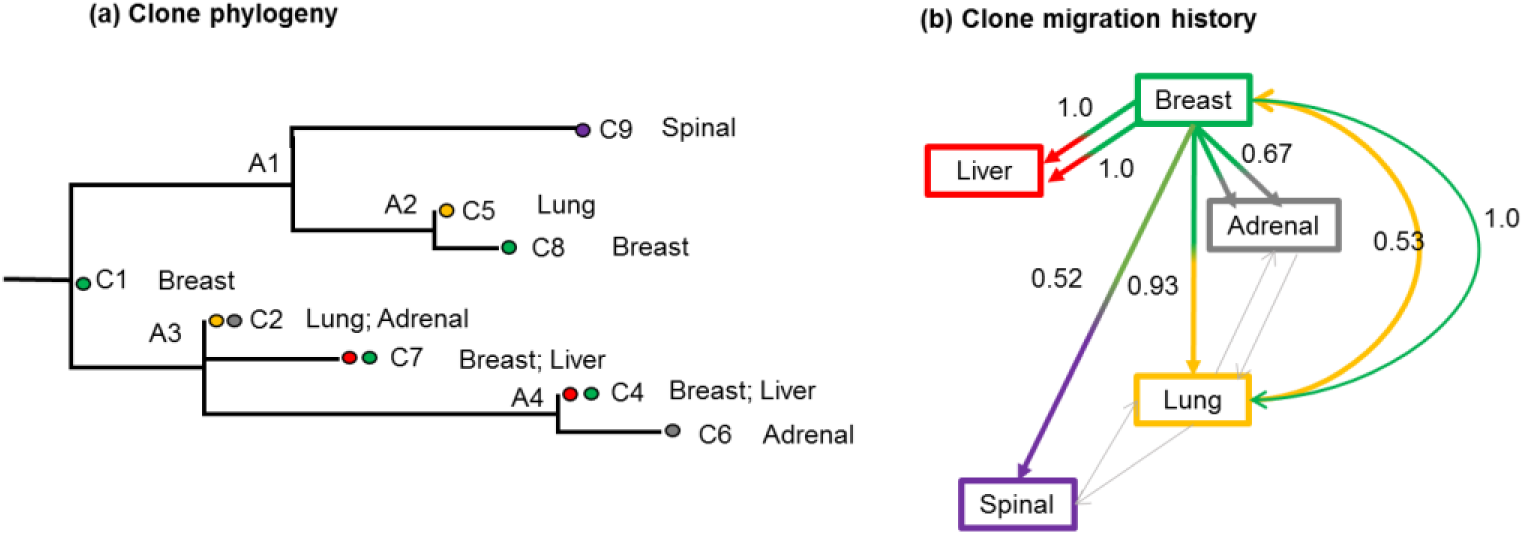
Patient A1 with basal-like breast cancer (Hoadley *et al*., 2016). (a) Clone phylogeny and tumor location of each clone reported in the original study. Nodes A1-A4 were ancestral nodes. (b) Clone migration history predicted by *PathFinder*. Inferences of migration paths included M→M and M→P paths. Predicted probabilities greater than 0.5 are depicted above the arrows. Colors correspond to the tumor location where clones were sampled from.

For example, the probability of the path to spinal was relatively low (0.52). The spinal tumor site contained only clone C9, and its direct ancestral node was A1. Although the ancestral tumor site of A1 was predicted to be breast (primary), all branches connected to this node A1 were relatively long with branch lengths similar to each other. Under this situation, the ancestral tumor site cannot be unambiguously determined by the information of branch lengths. As a result, the migration event that originated from the node A1 obtained low probability support, i.e., the P→M path (breast→spinal).

Similarly, the probabilities of migration paths to adrenal and the lung were not very high (0.67 and 0.53, respectively). In these cases, the ancestral tumor site at node A3 was challenging to infer by using only the phylogenetic information. The ancestral node A3 was leading to many clones that were found within the breast, while the lung and adrenal contained clone C2 that was directly connected to A3 with a zero branch length. Since node A3 was the direct descendant of the root of the phylogeny, the ancestral tumor site at the node A3 was likely breast. However, we cannot negate the possibility of lung or adrenal as the ancestral tumor sites at this node, resulting in low support for these migration paths to lung and adrenal sites. Interestingly, *PathFinder* detected a reseeding event from the lung tumor site (*P* = 1.0) (**Fig. 10b**). This is because clone C8, observed in the breast tumor, is a direct descendant clone of clone C5 that is found within the lung. Since clone C5 is not observed within any other tumor sites nor C5 has any other direct descendant clones, only a reseeding event can explain this observed pattern.

Overall, *PathFinder* predicted alternative migration histories for these two empirical datasets, including many seeding events between metastases as well as a reseeding event in which a metastatic clone moved back to the primary tumor. Our findings are supported by various studies in metastatic breast cancer that discuss extensive heterogeneity of tumors as a result of seeding or reseeding events by multiple clones between metastases (Yates *et al*., 2017; Savas *et al*., 2016). Developing *PathFinder* enabled us to discover more migration paths between clones and explore alternative migration histories.

## 4 Conclusions

Accurate computational methods for inferring cell migration routes are needed to answer fundamental questions in cancer biology, such as: How often do metastatic tumors arise from primary tumors (P→M) versus metastatic tumors (M→M)? How often do cells from metastases move back to primary tumors (reseeding M→P), and how often do tumors exchange clones (M↔M and P↔M)? We also need to know if these propensities differ among cancer types and patients. The sequencing of increasing numbers of cancer cells and tumors from many patients is poised to provide data essential to unravel the complexity of cancer cell movements. These data will not be able to fulfil their promise without the development of accurate methods to infer migration histories.

Therefore, the statistical estimation of clone migration histories is vital in cancer research as it can model the origin and movements of cancer cells between tumors. The only existent method for clone migration inferences is based on the maximum parsimony principle (El-Kebir *et al*., 2018), with attempts from researchers to also explore models borrowed from the field of biogeography (Alves *et al*., 2019; Chroni *et al*., 2019). We have presented a new Bayesian method that uses the clone phylogeny, including clone branch lengths, to estimate most likely migration histories. This approach increases the accuracy of estimated migration histories, and provides a direct way to compare alternative possible migration histories.

By analyzing the anatomy of errors in the simulated data, we have shown that many of the errors were caused by the lack of sufficient sampling of clones in each tumor site and a limited number of nucleotide variants for each tumor clone. This could be remedied by using clone-specific mutational signatures, structural variants, copy-number alterations, and epigenetic changes. We hope to integrate such information in the *PathFinder* approach to make it even more accurate, making it more useful as it is becoming easier to obtain genomic data from multiple tumors within a patient.

## Acknowledgments

We thank Allan George and Jiyeong Choi for their technical support.

## Author contributions

S.K. developed the original method, S.K. and S.M. refined the method and designed research; S.K., M.S., K.T., and S.M. implemented the algorithm; S.M., A.C., O.O., V.A., and T.V. performed the analysis; and S.K., S.M., and A.C. wrote the paper.

